# NF*ix*DB (Nitrogen Fixation DataBase) - A Comprehensive Integrated Database for Robust ‘Omics Analysis of Diazotrophs

**DOI:** 10.1101/2024.03.04.583350

**Authors:** Madeline Bellanger, Jose L. Figueroa, Lisa Tiemann, Maren L. Friesen, Richard Allen White

## Abstract

Biological nitrogen fixation is a fundamental biogeochemical process that transforms that provides fixed biologically available nitrogen by diazotrophic microbes. Diazotrophs anaerobically fix nitrogen using the nitrogenase enzyme which has three different gene clusters: 1) molybdenum nitrogenase (*nifDHK*) is the most abundant, followed by it’s alternatives 2) vanadium nitrogenase (*vnfDHK*), and 3) iron nitrogenase (*anfDHK*). Multiple databases have been constructed as resources for diazotrophic ‘omics analysis; however, an integrated database based on whole genome references does not exist. Here, we present NF*ix*DB (<u>N</u>itrogen <u>Fix</u>ation <u>D</u>ata<u>B</u>ase), a comprehensive integrated whole genome based database for diazotrophs, which includes all nitrogenases (*nifDHK*, *vnfDHK*, *anfDHK*) and nitrogenase-like enzymes (e.g., *nflDH*) linked to ribosomal operons (16S-5.8S-23S). NF*ix*DB was computed using Hidden Markov Models (HMMs) against the entire whole genome based Genome Taxonomy Database (GTDB R214), providing searchable reference HMMs for all nitrogenase and nitrogenase-like genes, complete ribosomal operons, both GTDB and NCBI/RefSeq taxonomy, and an SQL database for querying matches. We compared NF*ix*DB to *nifH* databases from Buckley, Zehr, Mise, and FunGene finding extensive evidence of *nifH*, in addition to *vnfH* and *nflH*. NF*ix*DB contains more than 4,000 verified *nifHDK* sequences contained on 50 unique phyla of bacteria and archaea. NF*ix*DB offers the first comprehensive nitrogenase database available to researchers.

## Introduction

Biological nitrogen fixation (BNF) is an ancient biogeochemical process on Earth, which is the conversion of atmospheric dinitrogen (N_2_) to fixed biologically available nitrogen as ammonium (NH_3_), completed by specialized microbes known as diazotrophs (Garcia et al., 2020). Nitrogen is essential to all life on the planet, and required for amino acid and nucleic acid synthesis, yet prior to the emergence of the enzyme nitrogenase, elemental nitrogen on the early Earth could only be fixed by lighting (Mancinelli and McKay, 1988). Prior to industrialization, the bioavailable nitrogen supplied by crop ratios required to support the ecosystem productivity was produced almost solely via biological nitrogen fixation by diazotrophs (Goyal et al., 2021).

For many decades it was thought that only symbiotic BNF, in which bacteria colonize specialized plant structures, provided significant amounts of ecosystem nitrogen, since high energy demands limit BNF to circumstances with adequate supplies of carbon (Goyal et al., 2021). In recent years, there has been a growing realization that free living nitrogen fixation (FLNF) can provide fixed nitrogen at rates equal to or greater than symbiotic nitrogen fixation and may be the dominant source of new nitrogen inputs to many terrestrial ecosystems (Van langenhove et al., 2020).

The nitrogenase enzyme as three variations which include different complex metal clusters that are oxygenic-sensitive metalloenzymes: 1) molybdenum-dependent nitrogenase (*nifDHK*) which is the most abundant, followed by 2) vanadium-dependent nitrogenase (*vnfDHK*), and 3) iron nitrogenase (*anfDHK*). All nitrogenases contain both catalytic and biosynthetic genes within the nitrogenase gene cluster. The catalytic genes all contain a dinitrogenase reductase, the *H* gene (e.g., *nifH*), which functions as an ATP-dependent electron donor, and a metalloenzyme heterotetramer of *D* and *K* genes, which are metal dependent iron protein alpha chain and iron protein beta chain (Burén et al., 2020). The biosynthetic gene cluster includes *nifB, nifE,* and *nifN*, which are required for FeMo-co biosynthesis, with *nifB, nifU, nifS, nifV*, and *nifM* are also required by the alternative nitrogenases (Burén et al. 2020). A subset of diazotrophs also contain alternative nitrogenases, *vnf* or *anf*. The *vnf* gene cluster contains the iron-vanadium cofactor, while the *anf* gene cluster contains the iron-iron cofactor (Schwartz et al., 2022). It is still widely debated which nitrogenase emerged first (Mus et al., 2019). Nitrogenase-like protoenzymes evolved first in methanogens for F_430_ cofactor biosynthesis and are known as Ni-sirohydrochlorin- a,c- diamide reductive cyclase (*nfl*) (Boyd et al., 2011; 2015). The *nfl* enzymes are ubiquitous in diazotrophic prokaryotes (Boyd et al., 2011; 2015). Bacteriochlorophyll (*Bch*) and Chlorophyll (*Chl*) biosynthesis (gene light-independent protochlorophyllide reductase) evolved from *nfl*, and are also nitrogenase-like homologs (related to cobalamin biosynthesis, F_430_ cofactor biosynthesis, and biosynthesis of chlorophyll and bacteriochlorophyll) (Staples et al., 2007).

A major gap in the current literature is the lack of a comprehensive database of nitrogenase enzymes that is rooted in whole-genomes, and thus, our ability to define diazotroph phylogeny or infer metabolic properties of genomes that are able to function effectively as diazotrophs is severely limited. The original nitrogenase databases were based on amplicon sequences that were not complete, due to high sequencing costs for whole genomes and lack of reference genomes (Gaby & Buckley, 2014, Heller et al., 2014). The most current *nifD* and *nifH* database from FunGene contains 19,514 *nifH* and 10,482 *nifD* sequences and alternative nitrogenases are not well defined (Fish et al., 2013). FunGene is currently no longer available as of 2022, as the website is no longer functional (Fish et al., 2013). The Buckley and Zehr lab groups have *nifH* specific databases publicly available on their groups’ websites, however they have not been updated since 2012 and 2017, respectively (Gaby & Buckley, 2014, Heller et al., 2014). The Mise lab group has classified *nifH* sequences and compiled this information into a database, but no information on alternative nitrogenases is available (Mise et al., 2021). To combat this, a novel database, NF*ix*DB (“<u>N</u>itrogen <u>Fix</u>ation <u>D</u>ata<u>B</u>ase”), was created. The inclusion of the *nifDK* genes and the alternative nitrogenases in a new database would be the first comprehensive collection of this data. Compiling sequences with connecting rRNA marker (16S-5.8S-23S) databases to nitrogenase genes via complete genomes will provide an extensive database that can be the foundation for the current and future studies of nitrogen fixation.

## Materials and Methods

An overview of the methods can be seen in **Figure 1**. Initial seed sequences for *nifDHK*, *anfDHK*, *vnfDHK*, *nflDH*, and *ChlBIN* were manually curated. The sequences were locally aligned using MAFFT (Katoh and Standley, 2013). An HMM of each seed sequence was created using HMMER’s hmmbuild (Eddy, 2020), then combined together to make a concatenated file of the *nifDHK*, *anfDHK*, and *vnfDHK* HMMs and a concatenated file of the *nflDH* and *ChlBIN* HMMs. Using HMMER’s hmmsearch (Eddy, 2020), each genome in the release 214 of the Genome Taxonomy Database (GTDB) (Parks et al., 2022) was examined for the presence of nitrogen fixation enzymes. Over 80,000 representative genomes were analyzed.

**Figure 1.**
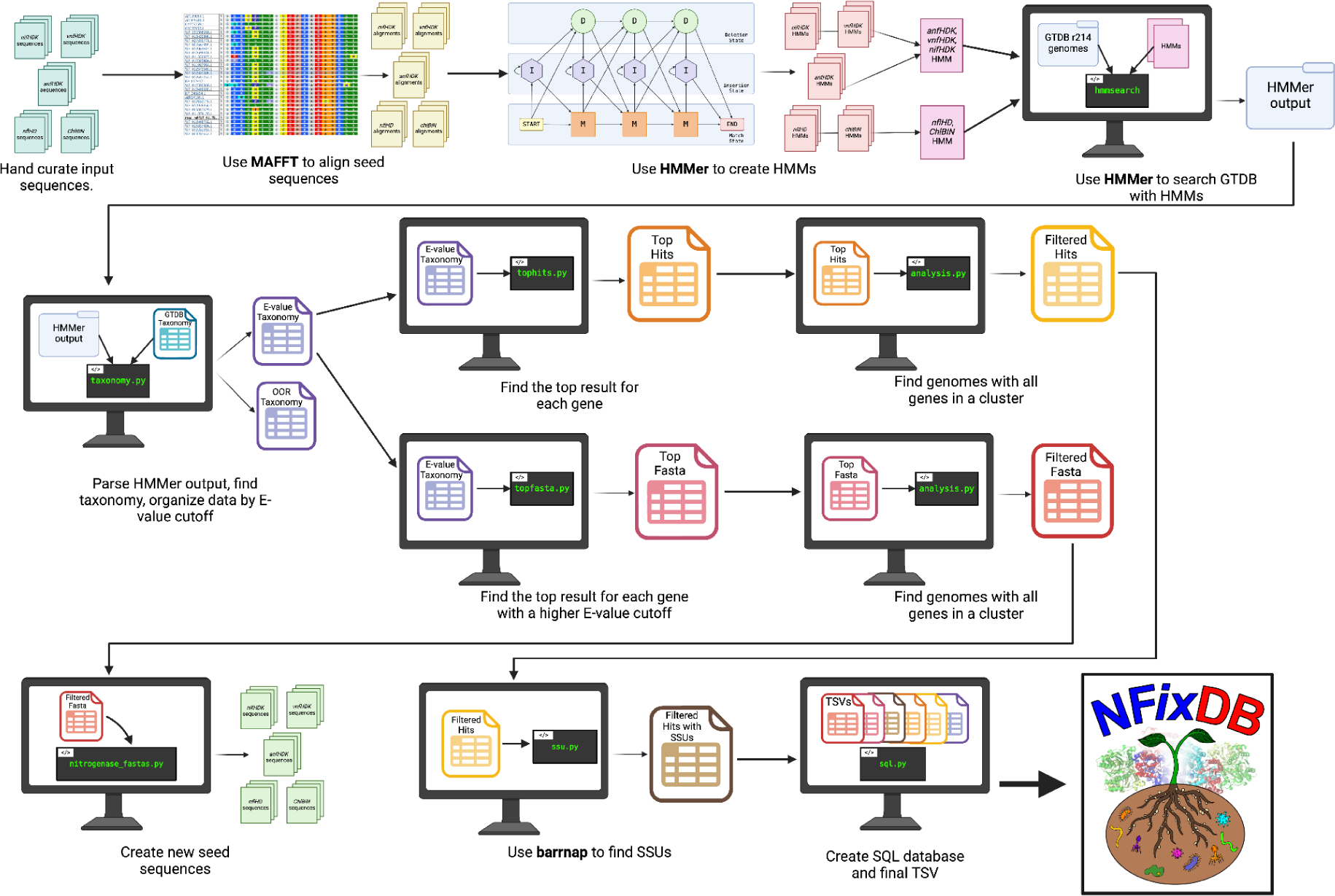
Flow graph of NF*ix*DB. Initial sequences were manually curated. Each gene was aligned using MAFFT. Then, an HMM was created with HMMer to search through all of the genomes in GTDB r214. The output was searched to find the best result for both the gene and the sequence ID. The sequences that occurred with all genes in a gene cluster were compiled together to make the new seed sequences.

The taxonomy of each protein sequence was found through both GTDB and the National Center for Biotechnology and Information (NCBI), with the NCBI taxonomy ID also being included. Additionally, an E-value cutoff was established at <9.9e^-10^. Any entry smaller than the cutoff value was placed into the evalue_taxonomy TSV file (**Zenodo**). From this TSV, the best result for each protein sequence was found and placed into the tophits TSV file (**Supplemental Table 6**). Additionally, another E-value cutoff of <9.9e^-15^, a bitscore cutoff value of >50, or an alignment length of >125 was established for updated seed sequence generation. The best result for each protein sequence was found (i.e., the result that scored the highest for both the gene and accession number). The sequences that fit within these parameters were placed into the topfasta TSV file (**Supplemental Table 6**). Both the tophits TSV and the topfasta TSV were examined to find genomes that contained all of the genes considered in the gene cluster. For instance, a genome would need to contain *nifH*, *nifD*, and *nifK* to pass this qualification. The tophits genomes that passed were placed into the filteredhits TSV and the topfasta genomes that passed were placed into the filteredfasta TSV (**Supplemental Table 6**).

The filteredfasta TSV was used to create an updated fasta file for each seed sequence. The ribosomal operons (16S-5.8S-23S) linked to each genome were identified using barrnap (Seemann, 2018). Multiple iterations were found to be unnecessary and led to bias towards similar genes (i.e. *nifH* mistaken for *vnfH*). The final database can be found in the NF*ix*DB TSV or as an SQL database, both on Zenodo, with each TSV mentioned being included as an SQL table. All genomes that were identified as containing nitrogenase, alternative nitrogenase, or pseudo-nitrogenase can be found on Zenodo.

Other *nifH* databases from Mise (Mise et al., 2021), Buckley (Gaby and Buckley, 2014), Zehr (Heller et al., 2014), and FunGene (Fish et al., 2013) were clustered using CD-HIT (Fu et al., 2012) at 100%, 99%, and 97% similarity. Representative sequences for clusters at 97% similarity were then analyzed using the HMMs that were used to curate NF*ix*DB to ensure our search results were as accurate as possible. Cutoffs for E-value <9.9e^-10^ and an alignment length >150 amino acids were put in place, with results being stored in the oDB class TSV (**Supplemental Table 7**). The best result for each sequence ID was identified and stored in the oDB hits TSV for all databases, along with a separate TSV for each individual database (**Supplemental Table 7**). As a secondary confirmation, global alignments were performed using SWORD (Vaser et al., 2016), with the representative sequences for clusters at 97% similarity as the queries and the final seed sequences from NF*ix*DB as the databases. This ensures that the final seed sequences produced from NF*ix*DB are accurate. Results that had an alignment length ≥220 amino acids, ≥50% identity, and E-value ≤9.9e^-10^ were analyzed to find the best result for each sequence ID. These entries were stored in separate TSVs for each database (**Supplemental Table 8**).

A significant increase in the number of sequences found in each gene cluster was observed from the initial seeds to the final seeds (**Supplemental Table 1, p < 0.01**). The seed sequences initially used were hand curated, making an uneven amount within each gene cluster. The production of the final seed sequences ensured an equal amount of sequences within each gene cluster (i.e., *nifH*, *nifD*, and *nifK* all have the same amount of sequences), and thus an equal amount in each new seed sequence FASTA.

## Results

Overall, NF*ix*DB resulted in the identification of more than 4,000 *nifHDK* genes, with an average length of 271 amino acids for *nifH*, 474 amino acids for *nifD*, and 472 amino acids for *nifK* (**Supplemental Figure 1**). Of all the sequences found to be from Proteobacteria, more than 50% were the *nifHDK* genes (**Figure 2**). Alphaproteobacteria and Deltaproteobacteria were the most common classes found in the *nifHDK* genes (**Figure 2**). In addition to the *nifHDK* genes identified, more than 250 *anfHDK* genes were found, with an average length of 270 amino acids for *anfH*, 505 amino acids for *anfD*, and 452 amino acids for *anfK* (**Supplemental Figure 1**). As with the *nifHDK* genes, the most common phyla found among the *anfHDK* genes were Proteobacteria (**Figure 2**). The most common classes in the *anfHDK* genes were Alphaproteobacteria and Gammaproteobactiera (**Figure 2**). Among the approximately 60 *vnfHDK* genes accurately identified, only three phyla were found (in order of most to least common): Proteobacteria, Cyanobacteria, and Firmicutes (**Figure 2**). Within those phyla, four classes were identified (in order of most to least common): Gammaproteobacteria, Alphaproteobacteria, Betaproteobacteria, and Bacilli (**Figure 2**). The lengths of *vnfH* averaged 270 amino acids, *vnfD* averaged 447 amino acids, and *vnfK* averaged 459 amino acids (**Supplemental Figure 1**).

**Figure 2.**
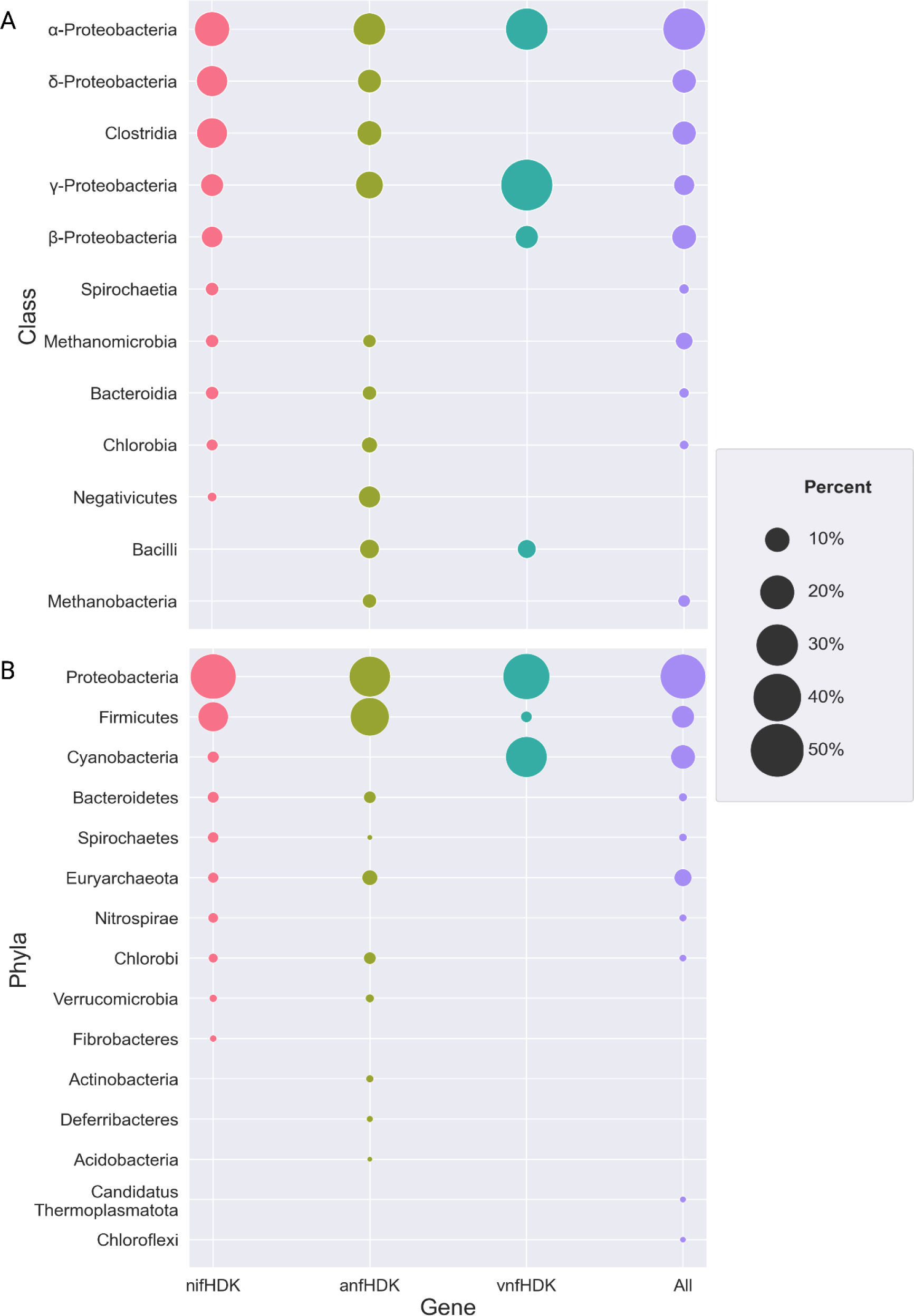
NF*ix*DB taxonomic classification at phyla and class level. The classes or phyla identified with the *nifHDK* genes are shown in pink. The classes or phyla identified with the *anfHDK* genes are shown in green. The classes or phyla identified with the *vnfHDK* genes are shown in blue. The top 10 overall classes or phyla are shown in purple. The size of each dot represents the percentage of occurrences among all of the classes or phyla identified. A) Top 10 classes for each nitrogenase gene group. B) Top 10 phyla for each nitrogenase gene group.

Of the 181 phyla analyzed, only 16% were found to have all the catalytic nitrogenase genes (*nifDHK* and/or the alternatives) (**Figure 2**). The majority of phyla with diazotrophs were Proteobacteria at more than 50%. Cyanobacteria held roughly 15% of organisms with diazotrophic activity (**Figure 2**).

We compared other *nifH* databases against NF*ix*DB, which include Buckley, Zehr, Mise, and FunGene. Prior to comparison we clustered to remove duplicates using CD-HIT (100%, 99%, and 97% similarity) then compared with HMMER3/HMMs and SWORD via global alignment. Clustering at 100% similarity resulted in a more than 50% decrease in the amount of sequences in both the Buckley and Zehr databases (**Figure 3, Supplemental Table 3, Supplemental Table 4**). The Mise and FunGene databases both had a less than 10% decrease per gene (**Figure 3, Supplemental Table 3, Supplemental Table 4**). Clustering down to 99% similarity resulted in a ∼60% decrease in the Buckley and Zehr databases (**Figure 3, Supplemental Table 3, Supplemental Table 4**). In the Mise and FunGene databases, the clusters resulted in a ∼20% decrease per gene (**Figure 3, Supplemental Table 3, Supplemental Table 4**). When clustering at 97%, the Buckley and Zehr databases decreased by ∼80%, leaving roughly 7,000 sequences in each database (**Figure 3, Supplemental Table 3, Supplemental Table 4**). The Mise and FunGene databases also saw decreases, ranging from 25% to 45% per gene (**Figure 3, Supplemental Table 3, Supplemental Table 4**).

**Figure 3.**
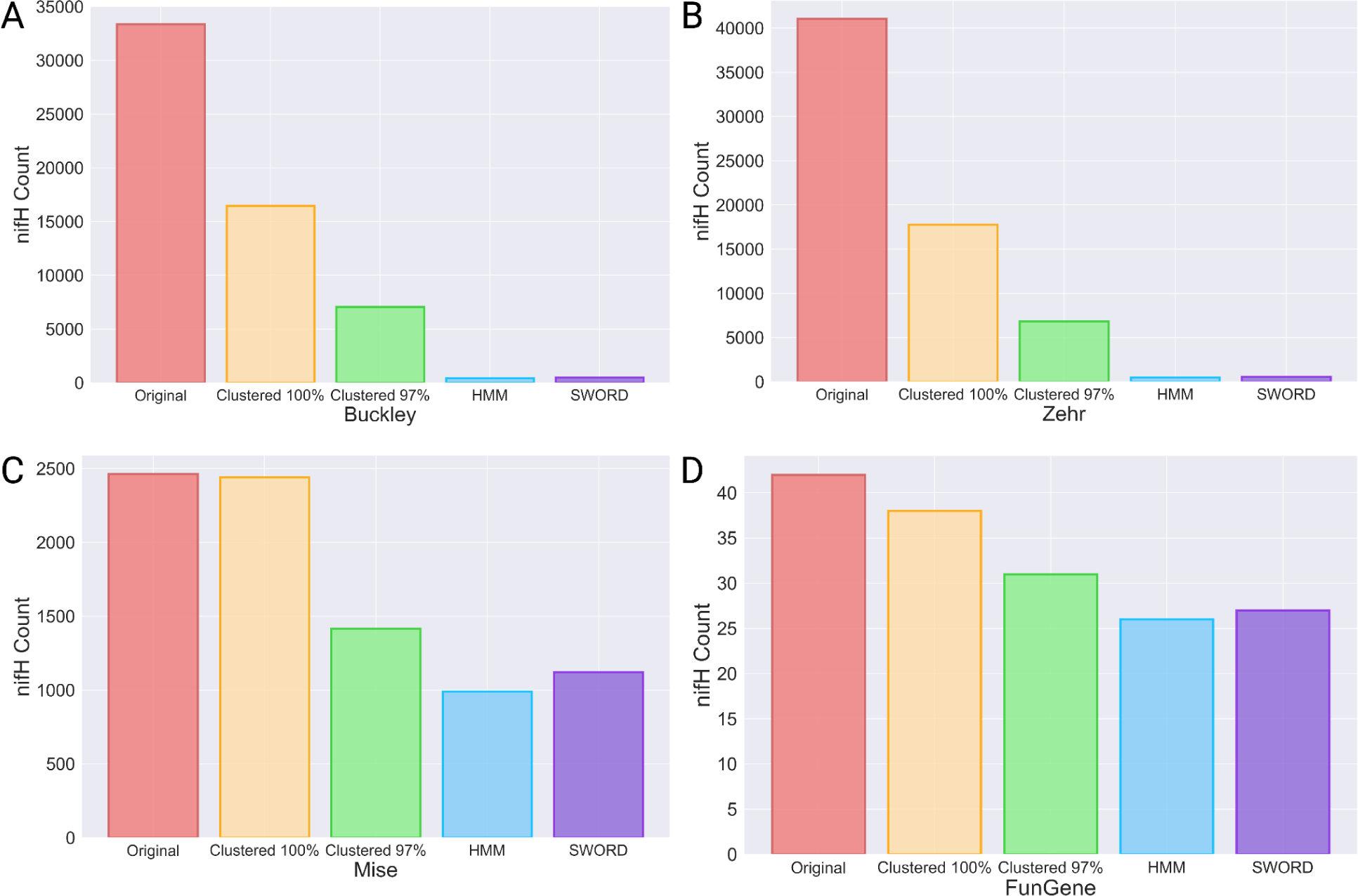
NF*ix*DB database comparison against Buckley, Zehr, Mise, and FunGene databases. The original count is shown in pink. Clustering at 100% similarity is shown in yellow and 97% similarity is shown in green. The count after classifying with our HMMs is shown in blue. The count after classifying with SWORD is shown in purple. A) Counts of sequences found in the Buckley database at each step. B) Counts of sequences found in the Zehr database at each step. C) Counts of sequences found in the Mise database at each step. D) Counts of sequences found in the FunGene database at each step.

**Figure 4.**
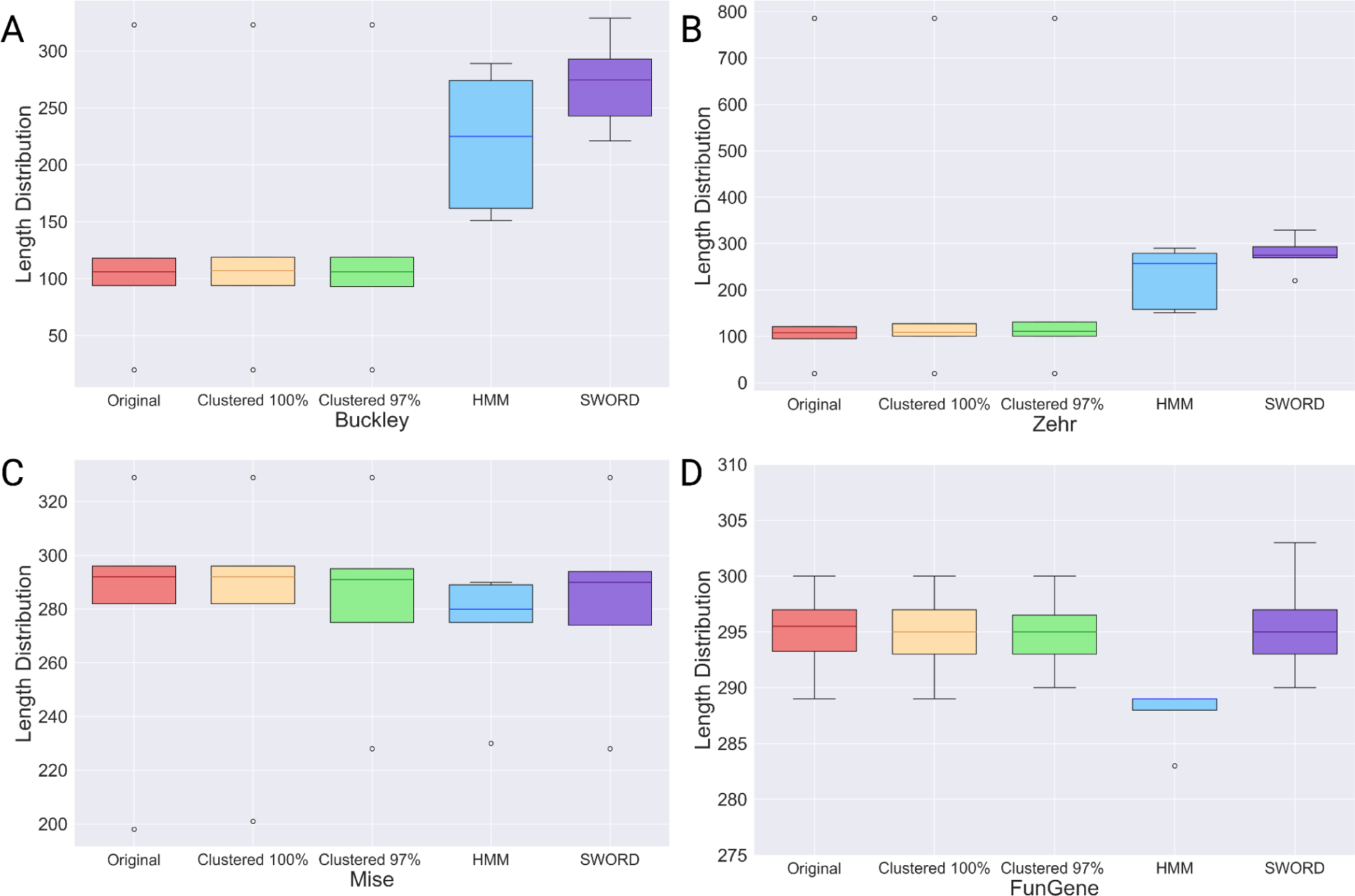
NF*ix*DB length distribution comparison against Buckley, Zehr, Mise, and FunGene databases. The original length distribution is shown in pink. Clustering at 100% similarity is shown in yellow and 97% similarity is shown in green. The length distribution after classifying with our HMMs is shown in blue. The length distribution after classifying with SWORD is shown in purple. A) Length distribution of sequences found in the Buckley database at each step. B) Length distribution of sequences found in the Zehr database at each step. C) Length distribution of sequences found in the Mise database at each step. D) Length distribution of sequences found in the FunGene database at each step.

When analyzing with both our HMMs and SWORD, these databases were found to have similar estimations of *nifH* sequences to what was originally estimated after clustering at 97% similarity (**Figure 3**, **Table 1**). The Mise database is split into three classifications of *nifH*. All three classifications were analyzed and 97% clustering resulted in 1,416 sequences total (**Supplemental Table 3**). Ten of those sequences did not pass our filtering, with an E-value <9.9e^-10^ and an alignment length >150 amino acids. Our HMMs revealed that the majority of sequences were found to be *nifH*. Roughly 30% of sequences analyzed were found to be *vnfH* or *nflH*. The SWORD analysis showed similar results, except fewer *vnfH* sequences and more *nifH* were identified (**Figure 3**, **Table 1**).

**Table 1.** NF*ix*DB database comparison summary table. Counts of the number of genes found in all of the outside databases after clustering at 97% similarity and classifying using both SWORD and HMMs. The gene that is shown in bold is the original gene identification from the corresponding database. All genes below that are genes that were identified through our classification.

| Database | HMM Analysis |  | Sword Analysis |  |
| --- | --- | --- | --- | --- |
| Original Gene |  |  |  |  |
| Detected Gene | Counts | % of Total | Counts | % of Total |
| **Buckley Total** | **627** |  | **606** |  |
| ***nifH*** | **627** |  | **606** |  |
| *anfH* | 34 | 5.42% | 33 | 5.45% |
| *ChII* | 2 | 0.32% | 2 | 0.33% |
| *nflH* | 54 | 8.61% | 41 | 6.77% |
| *nifH* | 444 | 70.81% | 500 | 82.51% |
| *vnfH* | 93 | 14.83% | 30 | 4.95% |
| **FunGene Total** | **241** |  | **241** |  |
| ***nifD*** | **190** |  | **190** |  |
| *nifD* | 187 | 98.42% | 190 | 100.00% |
| *vnfD* | 3 | 1.58% | 0 | N/A |
| ***nifH*** | **31** |  | **31** |  |
| *nifH* | 26 | 83.87% | 27 | 87.10% |
| *vnfH* | 5 | 16.13% | 4 | 12.90% |
| ***vnfD*** | **20** |  | **20** |  |
| *vnfD* | 20 | 100.00% | 20 | 100.00% |
| **Mise Total** | **1406** |  | **1406** |  |
| ***nifH*** | **1406** |  | **1406** |  |
| *nflH* | 113 | 8.04% | 111 | 7.89% |
| *nifH* | 990 | 70.41% | 1122 | 79.80% |
| *vnfH* | 303 | 21.55% | 173 | 12.30% |
| **Zehr Total** | **974** |  | **896** |  |
| ***nifH*** | **974** |  | **896** |  |
| *anfH* | 59 | 6.06% | 58 | 6.47% |
| *ChII* | 2 | 0.21% | 7 | 0.78% |
| *nflH* | 285 | 29.26% | 224 | 25.00% |
| *nifH* | 517 | 53.08% | 557 | 62.17% |
| *vnfH* | 111 | 11.40% | 50 | 5.58% |

The Zehr database clustered down to 6,831 sequences (**Supplemental Table 3**). Only 974 of those sequences passed our filters for HMM analysis, with an E-value <9.9e^-10^ and an alignment length >150 amino acids. Our HMMs revealed that more than 50% of the sequences analyzed were classified as *nifH*. Roughly 17% of the sequences were classified as *anfH* and *vnfH*. Nearly 30% of sequences were identified as *nflH* and a small subset of sequences were identified as *ChIl* (<1%). The SWORD analysis again showed similar results, with more *nifH* and less *nflH* and *vnfH* being identified over 896 sequences (**Figure 3**, **Table 1**).

The Buckley database was clustered down to 7,062 *nifH* sequences (**Supplemental Table 3**). Only 627 of those sequences passed our filters with an E-value <9.9e^-10^ and an alignment length >150 amino acids. More than 70% of those were classified as *nifH* in our analysis. Nearly 15% of sequences were classified as *vnfH*. The rest of the sequences were found to be *anfH* (∼5%), *nflH* (∼8%), and *ChIl* (∼0.3%). The SWORD analysis resulted in 606 sequence hits after filtering. Of those, more than 80% were identified as *nifH*. Nearly 5% were identified as *vnfH*. The rest were classified similar to the HMM results (**Figure 3**, **Table 1**).

FunGene clustered at 97% similarity led to 31 *nifH* seed sequences, 190 *nifD* seed sequences, and 20 *vnfD* seed sequences (**Supplemental Table 3**). It is important to note that FunGene does offer many more sequences for these genes, but does not consider all of those seed sequences. For the purpose of this study, only the seed sequences were analyzed. After our analysis with both HMMs and SWORD, it was found that all of the *vnfD* sequences are accurately classified. The majority of the *nifD* sequences were *nifD*, with ∼1.5% of them classified as *vnfD* when using HMMs. The *nifH* sequence results were similar using both HMMs and SWORD, with one more sequence being identified as *nifH* over *vnfH* when using SWORD. There were roughly 85% of sequences correctly identified as *nifH* and roughly 15% of sequences identified as *vnfH* (**Figure 3**, **Table 1**).

## Discussion

Generally, the measurement of *nifH* has been the gold standard for quantifying the diversity, abundance, presence, and potential activity of diazotrophs. In the era of highly cost effective next generation sequencing and high throughput quantitative PCR measurements, understanding the quality of the resulting *nifH* databases is critical to agriculture, food security, and bioenergy applications, which are all limited by nitrogen. Our analysis revealed that whole genome based curation provides a framework to further exploration of diazotrophs, highlighting the need to include alternative nitrogenases and nitrogenase-like genes.

Beginning with hand curated seed sequences, an alignment and an HMM was made for each gene. Every genome in GTDB was searched with each HMM to identify any potential nitrogenase genes. Further processing was done for each hit to ensure that only the top result for each accession number was kept. After finding genomes with all genes in a cluster present (i.e., *nifH*, *nifD*, and *nifK* present), new seed sequences were gathered to create NF*ix*DB.

The misclassification of *nifH* has had a large impact on the understanding of nitrogenase. In the past, alternative nitrogenases and pseudo-nitrogenases have not been classified well, despite *nifH*, *vnfH*, and *nflH* being very closely related. Throughout our database curation, *nifH* and *vnfH* were revealed to be continually mislabeled. For instance, *Azotobacter vinelandii DJ* (GCF_000021045.1) contains three genes labeled as *nifH*. One of these three genes is the true *nifH* (WP_012698831.1). The other two genes are *anfH* (WP_012703362.1) and *vnfH* (WP_012698955.1). Within *A. vinelandii DJ*, there are no annotated duplicates of a true *nifH*. Issues like this have led to *nifH* databases containing large amounts of *vnfH* and *nflH* (>10%).

Additionally, HMMs have a difficult time distinguishing between extremely closely related genes, like *nifH* and *vnfH*, however, a global alignment provides validation of the HMM results, when these methods are combined together. Recent advances in machine and deep learning such as convolutional neural networks (CNNs) could be applied to enhance detection of nitrogenases within genomes or metagenomic data.

There has been a lack of an all encompassing database for diazotrophs that contains alternative nitrogenases, which could lead to misinterpretation of nitrogenase diversity, presence, and activity within ecosystems. NF*ix*DB provides the first comprehensive whole genome based database for nitrogenase, alternative nitrogenases, and nitrogenase-like genes. Through NF*ix*DB, we provide a fundamental framework to unravel the diversity, presence, and potential activity of diazotrophs across the tree of life.

## Supplementary Materials

The following are available online at GitHub (https://github.com/raw-lab/NFixDB) and Zenodo (DOI: 10.5281/zenodo.10525001).

## Availability and implementation

NF*ix*DB scripts are written in Python and distributed under a BSD license. The source code and database of NFixDB is freely available at https://github.com/raw-lab/NFixDB.

## Data Availability Statement

Scripts, data, and SQL database are available on Zenodo (DOI: 10.5281/zenodo.10525001) and GitHub (https://github.com/raw-lab/NFixDB)

## Contributing to NFixDB and Fungene

NF*ix*DB as a community resource has recently acquired Fungene (Fish et al. 2013). We welcome contributions of other experts, expanding annotation of all domains of life (viruses, bacteria, archaea, eukaryotes). Please send us an issue on our NF*ix*DB GitHub

(https://github.com/raw-lab/NFixDB/issues). We will fully annotate your genome, add suggested pathways/metabolisms of interest, and make custom HMMs to be added to NF*ix*DB and FunGene. Also, NF*ix*DB is available within the metaomics tool MetaCerberus (https://github.com/raw-lab/MetaCerberus)

## Acknowledgments

We acknowledge the support of the following units of the University of North Carolina at Charlotte: the College of Computing and Informatics, the Bioinformatics Research Center, the Department of Bioinformatics and Genomics, Research and Economic Development, Academic Affairs, University Research Computing, and North Carolina Research Campus.

