## Supplemental for "NF*ix*DB (Nitrogen Fixation DataBase) - A Comprehensive Integrated Database for Robust ‘Omics Analysis of Diazotrophs"

All files mentioned in the manuscript can be found on GitHub (https://github.com/raw-lab/NFixDB) or Zenodo (DOI: 10.5281/zenodo.10525001).

| Seed Sequence | Initial Count | Final Count |
|---|---|---|
| *nifH* | 29 | |
| *nifD* | 29 | 4165 |
| *nifK* | 29 | |
| *anfH* | 16 | |
| *anfD* | 19 | 264 |
| *anfK* | 19 | |
| *vnfH* | 13 | |
| *vnfD* | 17 | 59 |
| *vnfK* | 16 | |
| *ChlN* | 18 | |
| *ChlI* | 18 | 3270 |
| *ChlB* | 18 | |
| *nflD* | 17 | 696 |
| *nflH* | 17 | |

**Supplemental Table 1.** Seed counts after database curation. While initial seed counts are different, the seed curation for the final seeds resulted in an equal amount for each gene cluster. This ensures that the seed sequences are accurate and recently active.

| Genes | E-value ≤ $e^{-10}$ | E-value ≤ $e^{-15}$, BS ≥ 50, aln length > 125 |
| --- | --- | --- |
| *nifHDK* | 4204 | 4165 |
| *vnfHDK* | 61 | 59 |
| *anfHDK* | 264 | 264 |
| *nflHD* | 715 | 696 |
| *ChlBIN* | 3338 | 3270 |
| *nifHDK, vnfHDK, anfHDK* | 3 | 3 |

**Supplemental Table 2**. Counts of each gene cluster after filtering with an E-value less than or equal to $e^{-10}$ or an E-value less than or equal to $e^{-15}$, a bitscore greater than or equal to 50, or an alignment length greater than 125 amino acids.

| Database | Gene | Original Count | 100% Cluster Count | 99% Cluster Count | 97% Cluster Count |
|---|---|---|---|---|---|
| Buckley | *nifH* | 33377 | 16476 | 13172 | 7062 |
| FunGene | *nifD* | 290 | 263 | 215 | 190 |
| | *nifH* | 42 | 38 | 35 | 31 |
| | *vnfD* | 32 | 32 | 25 | 20 |
| Mise | *nifH T1* | 1756 | 1743 | 1412 | 1011 |
| | *nifH T2* | 578 | 568 | 438 | 317 |
| | *nifH T3* | 131 | 131 | 107 | 88 |
| Zehr | *nifH* | 41074 | 17777 | 13620 | 6831 |

**Supplemental Table 3.** Counts of representative sequences from outside databases after clustering with CD-HIT at 100%, 99%, and 97% similarity.

| Database | Gene | 100% Count Decrease | 99% Count Decrease | 97% Count Decrease |
|---|---|---|---|---|
| Buckley | *nifH* | 50.64% | 60.54% | 78.84% |
| FunGene | *nifD* | 9.31% | 25.86% | 34.48% |
| | *nifH* | 9.52% | 16.67% | 26.19% |
| | *vnfD* | 0.00% | 21.88% | 37.50% |
| Mise | *nifH T1* | 0.74% | 19.59% | 42.43% |
| | *nifH T2* | 1.73% | 24.22% | 45.16% |
| | *nifH T3* | 0.00% | 18.32% | 32.82% |
| Zehr | *nifH* | 56.72% | 66.84% | 83.37% |

**Supplemental Table 4.** Percentage decrease of representative sequences from outside databases after clustering with CD-HIT at 100%, 99%, and 97% similarity.

| Gene | Buckely | FunGene | Mise | Zehr |
|---|---|---|---|---|
| Original Total | 33377 | 364 | 2465 | 41074 |
| 97% Total | 7062 | 241 | 1416 | 6831 |
| *anfH* | 34 | 0 | 0 | 59 |
| *nifD* | 0 | 187 | 0 | 0 |
| *nifH* | 444 | 26 | 990 | 517 |
| *vnfD* | 0 | 23 | 0 | 0 |
| *vnfH* | 93 | 5 | 303 | 111 |
| *nflH* | 54 | 0 | 113 | 285 |
| *ChlI* | 2 | 0 | 0 | 2 |
| **Totals** | **627** | **241** | **1406** | **974** |

**Supplemental Table 5.** Comparison of FunGene, Mise, Buckley, and Zehr databases for *nifH* gene content. The counts of each gene were identified after clustering the databases at 97% similarity with CD-HIT and analyzed using our HMMs. It should be noted that FunGene has 31 *nifH* seeds, 190 *nifD* seeds, and 20 *vnfD* seeds, post clustering. The rest of the databases are entirely *nifH*.

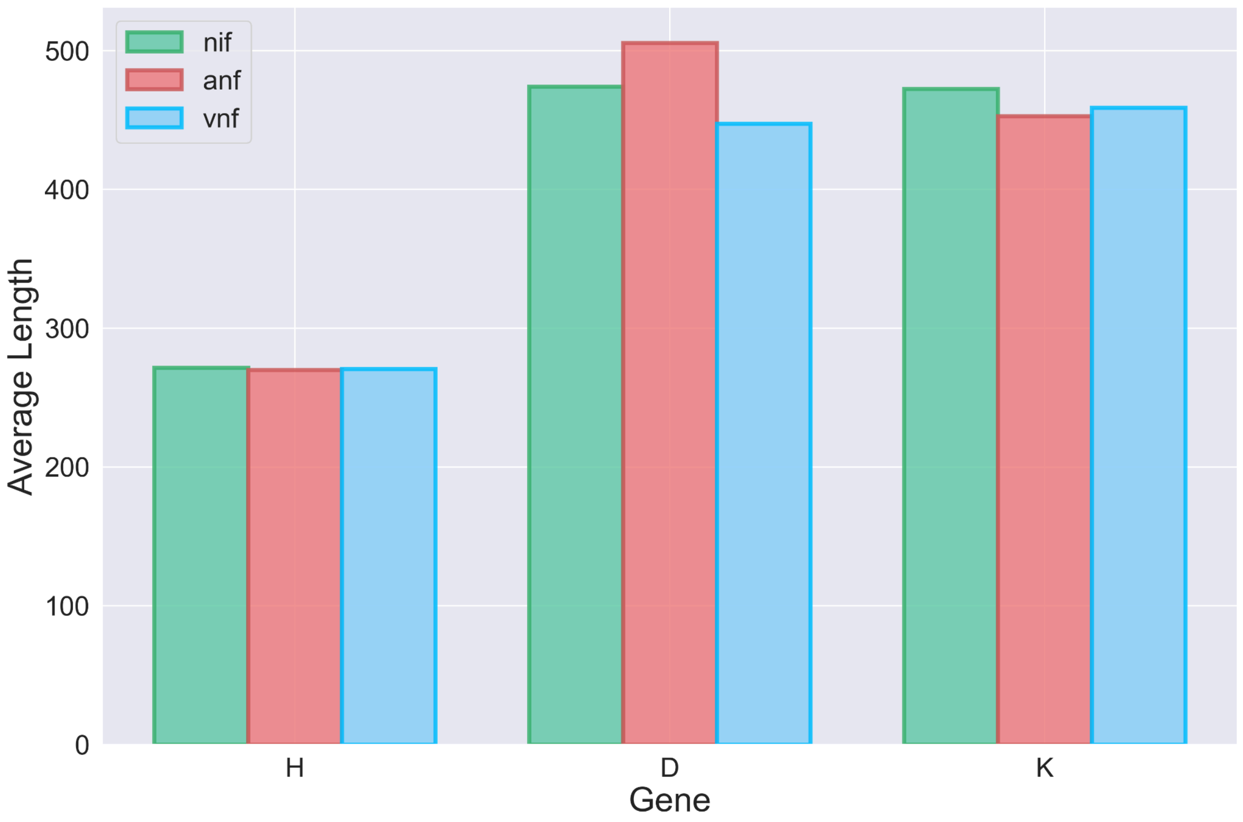


**Supplemental Figure 1.** Average length of all nitrogenase genes in NFixDB. Genes are grouped together by *H* (*nifH*, *anfH*, *vnfH*), *D* (*nifD*, *anfD*, *vnfD*), and *K* (*nifK*, *anfK*, *vnfK*). Green bars represent the *nif* genes. Pink bars represent the *anf* genes. Blue bars represent the *vnf* genes.

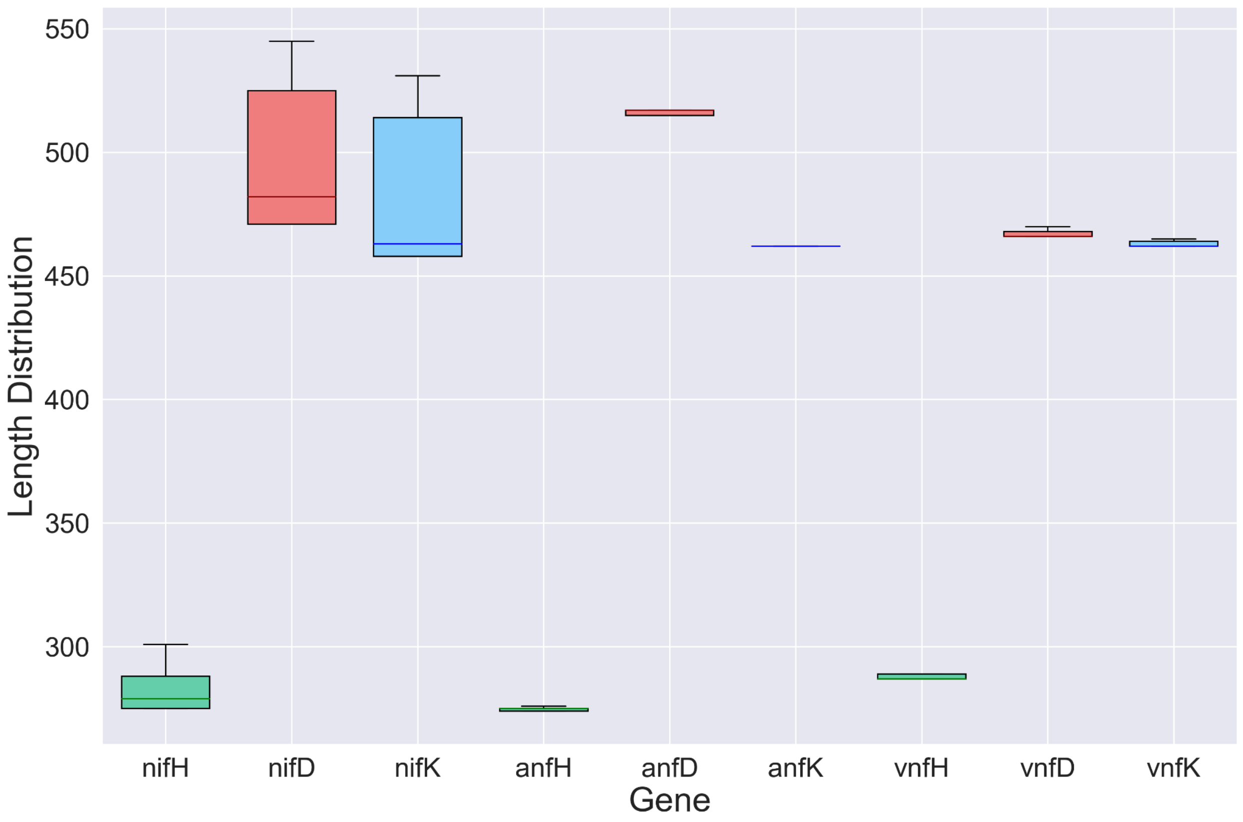


**Supplemental Figure 2.** Length distribution among each nitrogenase gene group. Genes are grouped together by *nif* (*nifH*, *nifD*, *nifK*), *anf* (*anfH*, *anfD*, *anfK*), and *vnf* (*vnfH*, *vnfD*, *vnfK*). Green bars represent the *H* genes. Pink bars represent the *D* genes. Blue bars represent the *K* genes.

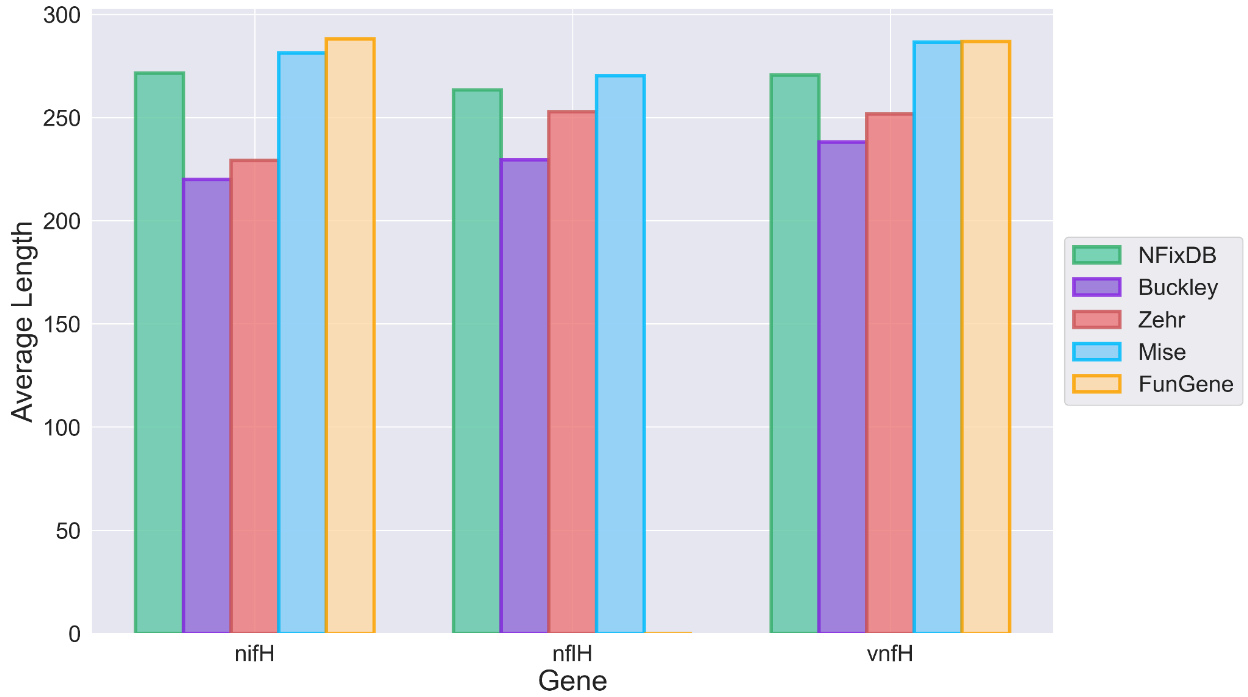


**Supplemental Figure 3.** Average length of *nifH*, *nflH*, and *vnfH* in each database after clustering at 97% and classifying using our HMMs. Our classification instituted a cutoff of an E-value <$9.9e^{-10}$ and an alignment length >150 amino acids. FunGene had no sequences identified as *nflH*.

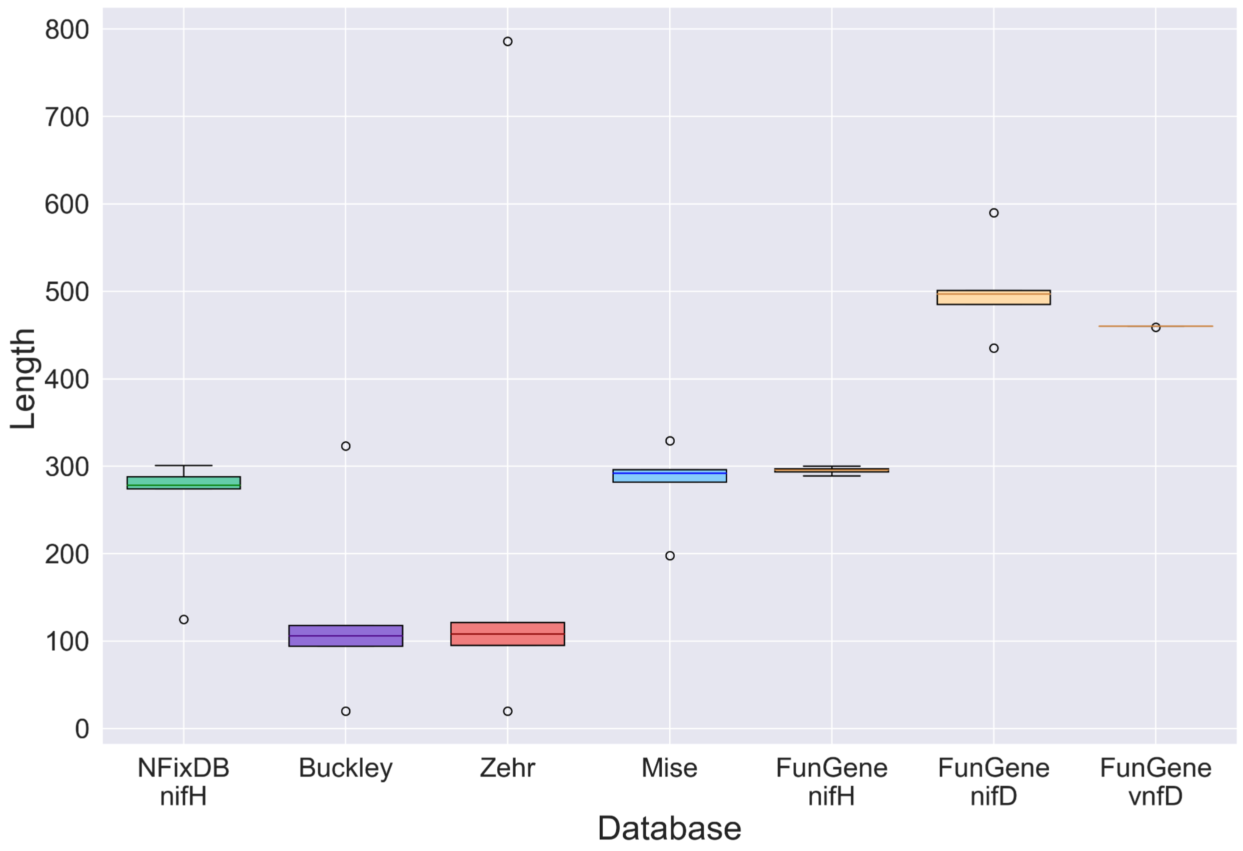


**Supplemental Figure 4.** Length distribution of sequences found in the databases before clustering or classification in comparison to the *nifH* length distribution found in NF*ix*DB. The Buckley, Zehr, and Mise databases all contain *nifH*. The FunGene database contains *nifH*, *nifD*, and *vnfD*.

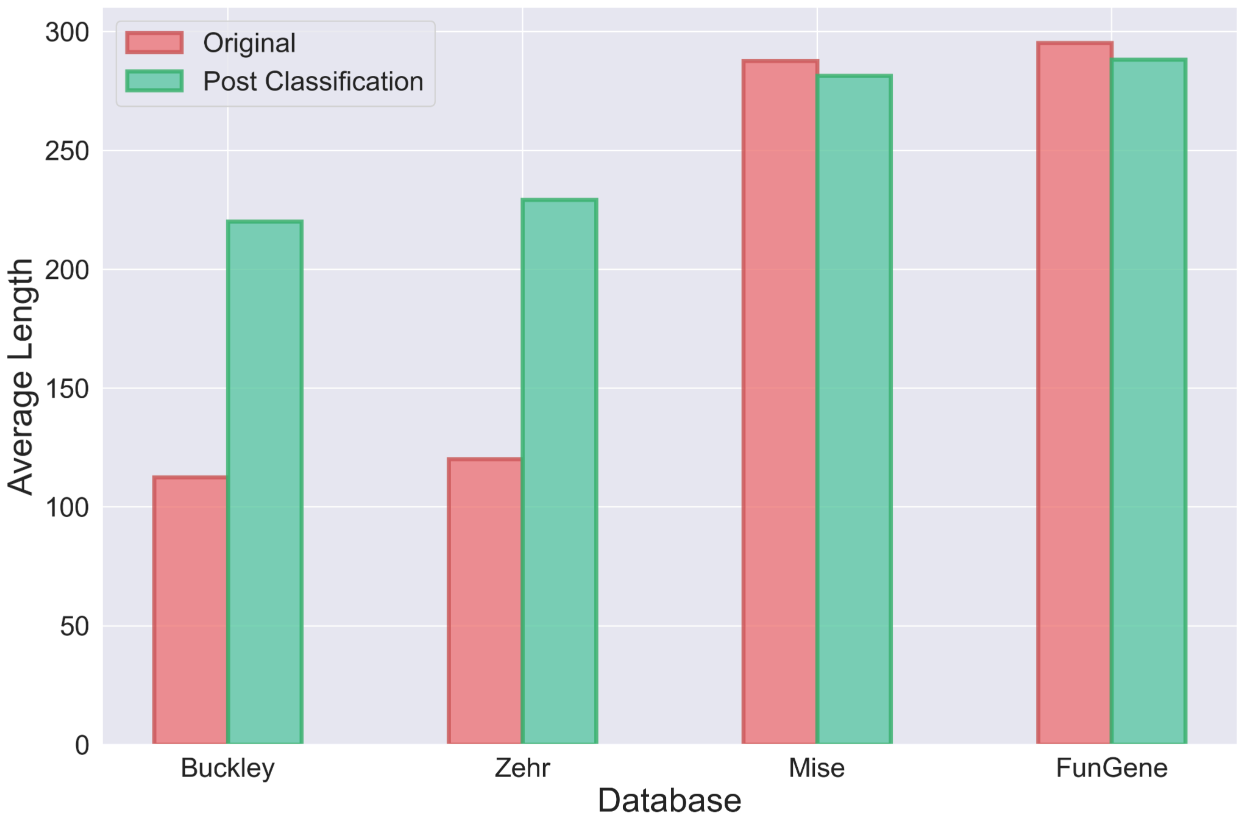


**Supplemental Figure 5.** Average length of *nifH* sequences in the original database compared to the *nifH* length in the classified database. Classification includes clustering at 97% similarity with CD-HIT, searching with our HMMs, and results with an E-value $<9.9e^{-10}$ and an alignment length >150 amino acids. The original average length is shown in pink and the classified average length is shown in green.

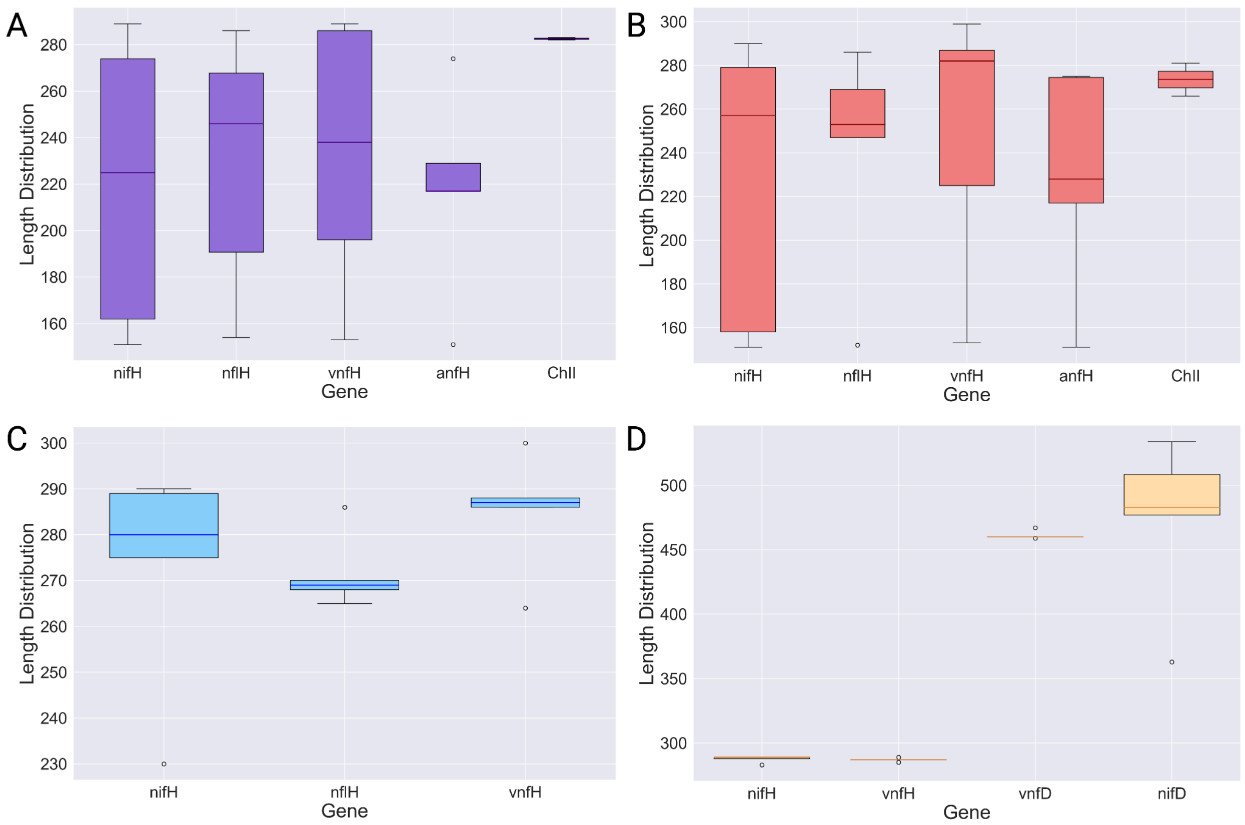


**Supplemental Figure 6.** Length distribution of each gene found in the various databases after clustering at 97% and classifying using our HMMs, with a cutoff of E-value <$9.9e^{-10}$ and alignment length >150 amino acids. A) Length distribution of genes identified in the Buckley database, shown in purple. Genes included are *nifH*, *nlfH*, *vnfH*, *anfH*, and *ChlI*. B) Length distribution of genes identified in the Zehr database, shown in pink. Genes included are *nifH*, *nlfH*, *vnfH*, *anfH*, and *ChlI*. C) Length distribution of genes identified in the Mise database, shown in blue. Genes included are *nifH*, *nlfH*, and *vnfH*. D) Length distribution of genes identified in the FunGene database, shown in yellow. Genes included are *nifH*, *vnfH*, *vnfD*, and *nifD*.

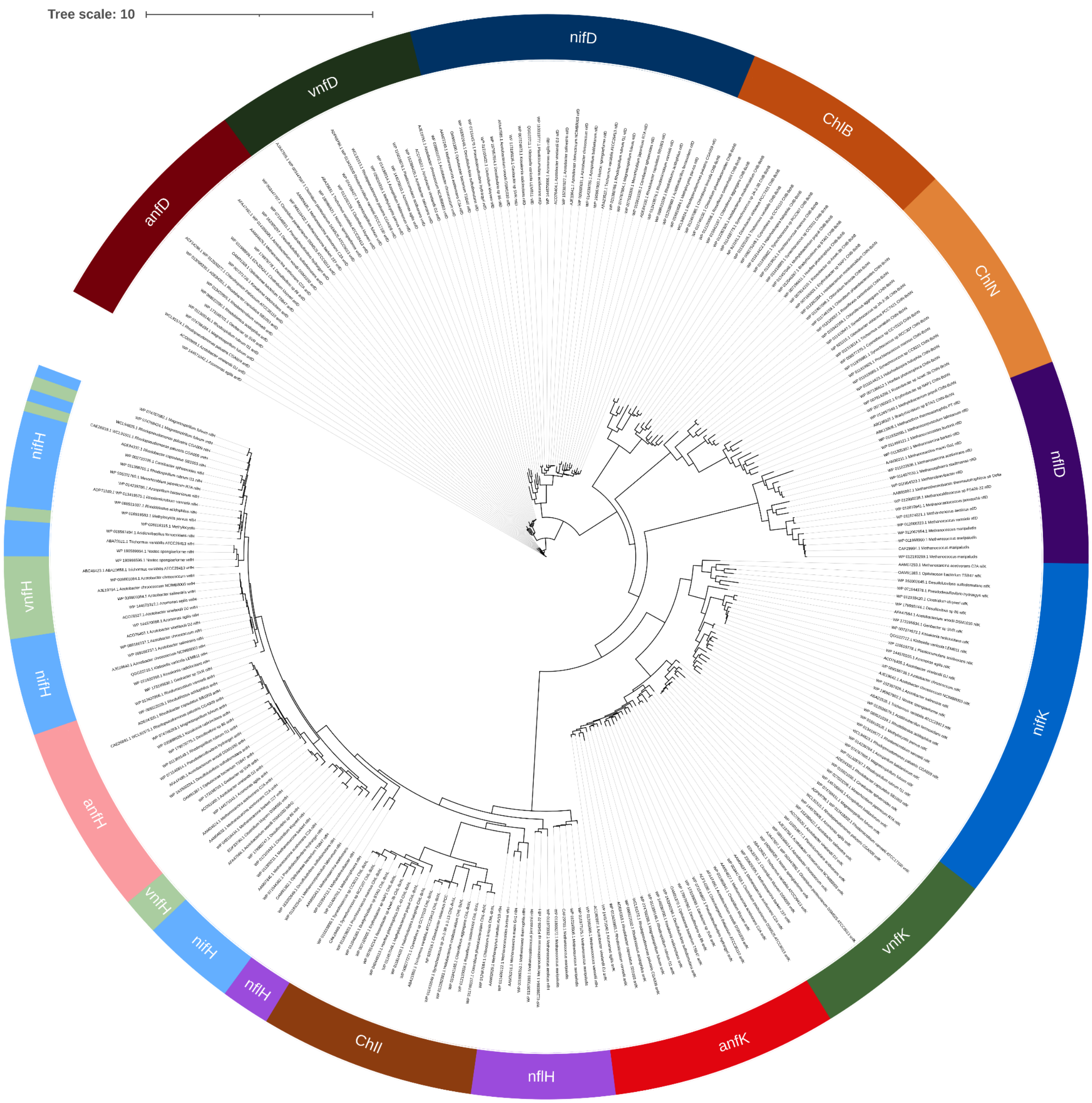


**Supplemental Figure 7.** Phylogenetic tree of initial, hand curated, seed sequences. The H genes, specifically *nifH* and *vnfH*, are phylogenetically ambiguous, hence the intertwining of *nifH*, *anfH*, and *vnfH*. The *anf* genes are shown in shades of red. The *vnf* genes are shown in shades of green. The *nif* genes are shown in shades of blue. The *Chl* genes are shown in shades of orange. The *nfl* genes are shown in shades of purple.
